# Genomic and epigenomic mapping of leptin-responsive neuronal populations involved in body weight regulation

**DOI:** 10.1101/601070

**Authors:** Fumitaka Inoue, Walter L. Eckalbar, Yi Wang, Karl K. Murphy, Navneet Matharu, Christian Vaisse, Nadav Ahituv

**Affiliations:** Department of Bioengineering and Therapeutic Sciences, University of California San Francisco, San Francisco, California, 94158, USA; Institute for Human Genetics, University of California San Francisco, San Francisco, California, 94158, USA; Diabetes Center, University of California San Francisco, San Francisco, California, 94143, USA

## Abstract

Genome wide association studies (GWAS) in obesity have identified a large number of noncoding loci located near genes expressed in the central nervous system. However, due to the difficulties in isolating and characterizing specific neuronal subpopulations, few obesity-associated SNPs have been functionally characterized. Leptin responsive neurons in the hypothalamus are essential in controlling energy homeostasis and body weight. Here, we combine FACS-sorting of leptin-responsive hypothalamic neuron nuclei with genomic and epigenomic approaches (RNA-seq, ChIP-seq, ATAC-seq) to generate a comprehensive map of leptin-response specific regulatory elements, several of which overlap obesity-associated GWAS variants. We demonstrate the usefulness of our leptin-response neuron regulome, by functionally characterizing a novel enhancer near Socs3, a leptin response-associated transcription factor. We envision our data to serve as a useful resource and a blueprint for functionally characterizing obesity-associated SNPs in the hypothalamus.

## Introduction

Obesity dramatically increases the risk of morbidity from hypertension, dyslipidaemia, diabetes and cardiovascular diseases and is a major public health concern^1–3^. Both environmental and genetic factors are involved in the onset and progression of weight gain^4^, with genetic factors being the major component^5–7^. By building on major advances in the description of molecular pathways implicated in food intake regulation and the control of body weight in mice, a number of rare obesity causing protein-coding variants have been identified in humans^8^. Most of the genes implicated in these monogenic forms of obesity encode proteins of the leptin axis and brain expressed targets of leptin in the hypothalamus^9,10^.

In parallel to monogenic causes of obesity, genome-wide association studies (GWAS) have identified over 250 loci for body mass index (BMI)^10^. Most of these GWAS loci represent clusters of common SNPs in noncoding regions^11^ that likely alter gene expression and have been found to reside primarily near genes involved in central nervous system (CNS)-related processes^12^. As the associated noncoding variant/s are not necessarily the causative variant/s, pinpointing the relationship between variants and causal regulatory elements and their target genes remains challenging. Gene regulatory maps, such as those generated by the ENCODE project, can allow the detection of potentially causative sequences nearby GWAS hits^13–15^. However, as gene regulatory elements, such as enhancers, tend to be cell type-specific^16^ and the hypothalamus contains a complex set of neuronal subtypes, the ability to identify obesity-associated regulatory elements remains highly constrained.

Here, we demonstrate the feasibility of establishing a comprehensive map of activated DNA regions in a sub-population of hypothalamic neurons by focusing on first order leptin-responsive neurons. We show that this map is consistent with the transcriptional activity of these neurons, leads to identification of relevant enhancers and provides a physical blueprint for identifying and testing the relevance of human obesity associated SNPs uncovered by GWAS.

## Results

### Nuclei isolation from leptin-responsive neurons

The hypothalamus is a complex brain structure composed of neuronal and non-neuronal cells such as astrocytes, microglia, oligodendrocyte and endothelial cells^17^. This complexity also makes separating these cells from one another for subsequent genomic analyses extremely complex. To map the genes and regulatory elements that are active in leptin-responsive neurons, a subset of hypothalamic neurons that includes Pomc, Agrp/Npy neurons, we generated mice in which the nuclei of these cells was fluorescently labeled and used these nuclei for subsequent cell isolation. Specifically, we crossed *Leprb^cre^* mice that carry an internal ribosome entry site (IRES) followed by CRE recombinase downstream of exon 18b of the *Lepr* gene^18^, with mice that express, in a Cre dependent manner, the SUN1 nuclear membrane protein fused to GFP^19^ (**Fig. 1a**). We confirmed that GFP labelling was restricted to the nucleus of subpopulation of neurons in the Arcuate (ARC), dorso-(DMH) and ventromedial hypothalamus (VMH), recapitulating endogenous *Leprb* expression pattern (**Fig. 1b**). We then established a protocol to isolate the cell nuclei from dissected hypothalamus (see Methods). This was followed by FACS sorting to separate the GFP positive from the GFP negative nuclei (**Fig. 1c, Supplementary Figure 1**). As the nutritional and/or the leptin-activation status may affect the transcriptome or regulome of these neurons, we performed the nuclear isolation under different conditions. Groups of mice were either fed, fasted or (to isolate the specific effect of leptin) given leptin injections chronically or acutely during fasting and experiments were carried out at the same time points during the day (**Fig. 1d**). For each condition, nuclei isolations were done from hypothalamus pooled from both male and female (to account for sex-specific differences) adult mice (2-4 months old) (**Supplementary Table 1**). For each assay, at least 5 mice per RNA-seq or ATAC-seq and at least 20 mice for ChIP-seq with three biological replicates for RNA-seq (total of 60 mice) and two for ATAC-seq (total of 40 mice) and ChIP-seq (total of 160 mice) were used.

**Fig. 1.**
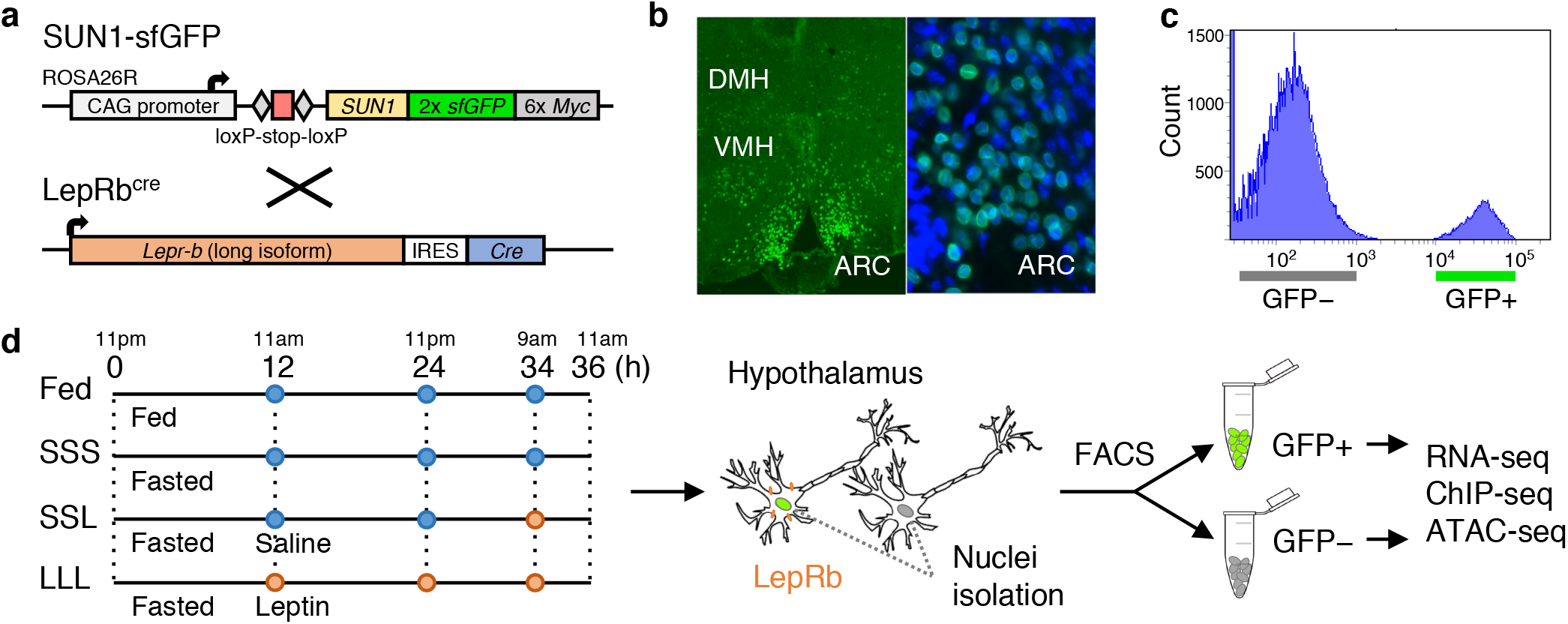
Isolation of LepRb GFP-positive nuclei. **a** *R26-CAG-LSL-Sun1-sfGFP-myc* homozygous mice were crossed with *LepRb^cre^* mice to obtain mice that express sfGFP in LepRb expressing cell nuclei. **b** Immunostaining showing sfGFP expression in the dorsomedial hypothalamus (DMH), ventromedial hypothalamus (VMH) and arcuate nucleus (ARC). On the right panel is a zoom in on the ARC along with DAPI staining (blue). **c** FACS histogram showing clear separation of GFP-positive from GFP-negative nuclei obtained from *LepRb^cre^* X *R26-CAG-LSL-Sun1-sfGFP-myc* heterozygous mice. Similar results were obtained in all the experimental groups (**Supplementary Figure 1**). **d** *LepRb^cre^* X *R26-CAG-LSL-Sun1-sfGFP-myc* heterozygous mice were fed or fasted for 36 hours, during which saline (blue) or leptin (orange) were injected at various time points. Cell nuclei were isolated and sorted via FACS, followed by RNA-seq, ChIP-seq, and ATAC-seq.

### Nuclear transcriptome of leptin-responsive neurons

In order to validate enrichment for Leprb expressing neuron nuclei, prior to mapping regulatory elements by H3K27ac ChIP-seq and ATAC-seq, and to provide a solid basis for matching the transcriptional status with the gene regulatory status in nuclei from leptin-responsive neurons, we analyzed the transcriptome of the sorted nuclei obtained by our method. We performed RNA-seq both on GFP-positive and GFP-negative sorted nuclei in all physiological conditions tested. The specificity of our nuclei isolation protocol was validated by the high level of enrichment of *LepR* and by the strong depletion of genes specifically expressed in oligodendrocytes (*Mag*), microglia (*Cx3cr1*) and astrocytes (*Aldh1l1*) (**Fig. 2a**). Comparison of our isolated GFP positive nuclei population versus all other cell types (GFP negative) found 1,897 genes to be significantly enriched in this population with *Lepr* being the most differentially expressed gene (**Fig. 2b**). Gene ontology analysis for genes that are differentially expressed in GFP-positive sorted nuclei showed enrichment for genes that function in energy homeostasis, such as eating behavior and regulation of appetite (**Supplementary Figure 2a**). Weighted correlation network analysis (WGCNA^20^), an unsupervised method to identify modules of co-expressed genes, found 10 modules to be highly correlated with GFP-positive and GFP-negative cell states (**Supplementary Figure 3a**). Gene ontology analysis of two of the most significantly correlated modules, ME3 and ME6, showed an enrichment for genes involved in neuronal activity for ME3 and a variety of terms for ME6 (**Supplementary Figure 3b**).

**Fig 2.**
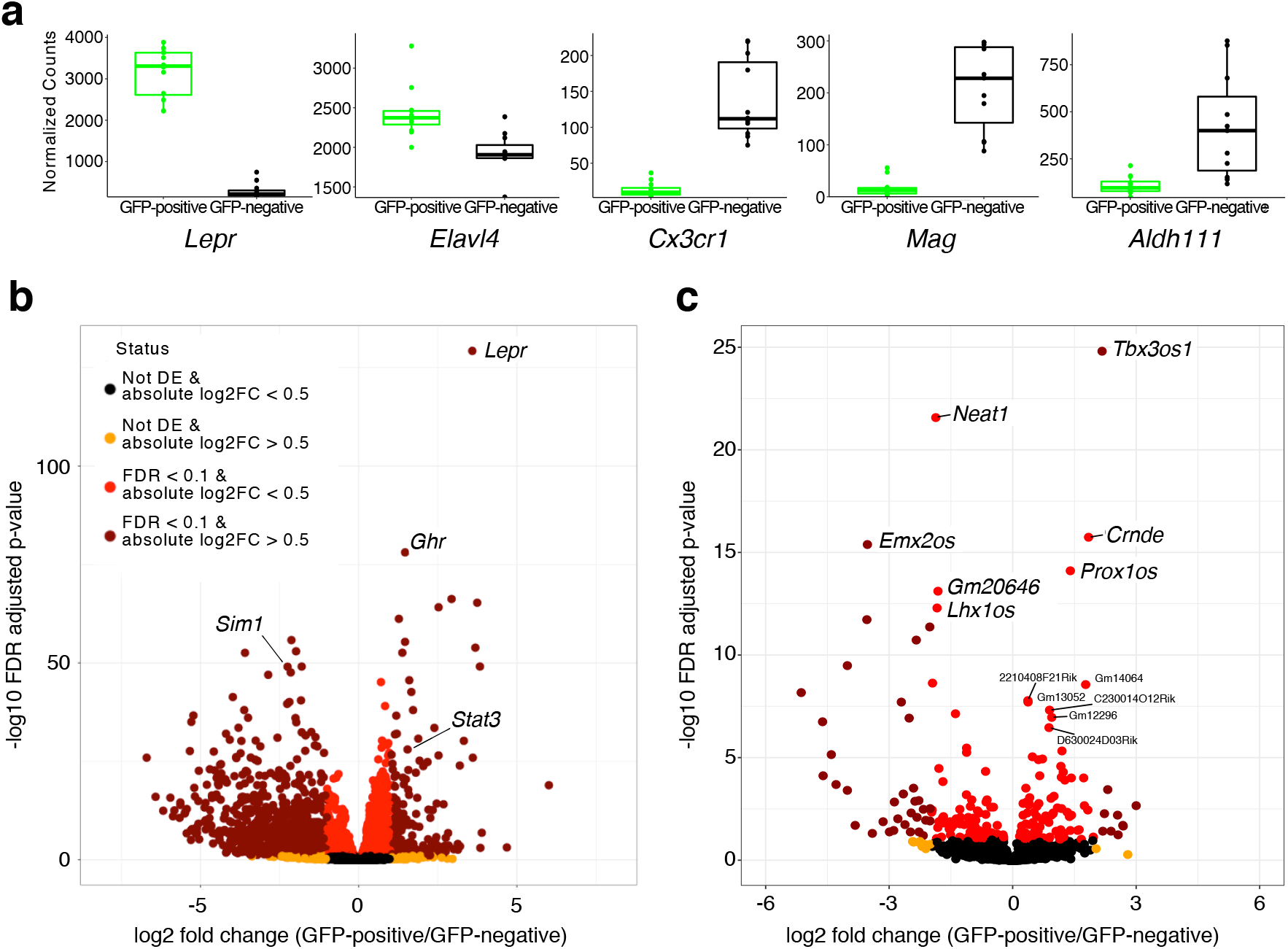
LepRb-GFP positive nuclei transcriptome. **a** Comparison of normalized counts for *Lepr* and genes specific for neurons (Elavl4(HUD), microglia (Cx3cr1), oligodendrocytes (Mag) and astrocytes (Adh1l1) in transcriptomes from GFP positive and GFP negative nuclei. Box plots represent the 25th to 75th percentiles and the midline indicates the median. Each point represents data from five pooled hypothalamus. **b** Volcano plot showing all differentially expressed genes when comparing LepRb GFP positive to GFP negative nuclei. **c** Volcano plot showing all differentially expressed lncRNAs when comparing LepRb GFP positive to GFP negative nuclei. Selected genes and lncRNAs are labeled. n=12 biologically independent replicates while pooling all physiological conditions tested (**a-c**). P-values given from Wald test and correct for multiple testing using the Benjamini & Hochberg procedure (FDR; **b** and **c**).

In addition to a global analysis, we also looked at individual genes. We detected significantly higher levels of expression in GFP-positive sorted nuclei of other leptin pathway associated genes such as *Stat3* (**Fig. 2b, Supplementary data file 1**). In addition, we observed that the growth hormone receptor (*Ghr*) gene, that was recently reported to be co-expressed with *Lepr* in the hypothalamus and plays an important role in glucose metabolism^21^, is also significantly enriched in the leptin responsive transcriptome.

While the difficulty of isolating whole neurons from adult mice has restricted the capacity to perform transcriptome analysis on neuronal sub-populations, Translating Ribosome Affinity Purification (TRAP), a method in which genetically labeled ribosomes can be immunoprecipitated from neuronal homogenates, has recently allowed the identification of translated transcripts (translatome) from specific neurons, including leptin-responsive neurons^22,23^. Comparison of our results to the previously published TRAP-seq dataset that analyzed the LepRb-neuron transcriptome^22^, found 88 upregulated (37.4%) and 175 downregulated (65.8%) genes that also showed up and down regulation in LepRb positive and LepRb negative cells in our dataset (**Supplementary Figure 4a**). In addition, recent work using TRAP-seq has implicated activating transcription factor 3 (*Atf3*) to be increased in LepRb neurons and function as a mediator of leptin signaling^23^. Analyses of our data also shows *Atf3* to be upregulated in LepRb neurons (**Supplementary Figure 4b**). Importantly, unlike translatome analysis, nuclear transcriptomes also allows for the analysis of non-translated RNA species. We therefore analyzed our RNA-seq dataset for long noncoding RNAs (lncRNAs) that are differentially expressed compared to all other cell types (GFP negative). We found numerous unique and 79 differentially expressed lncRNAs (33 increased and 46 decreased) (**Fig. 2c, Supplementary data file 1**).

To identify genes that specifically respond to physiological changes in leptin-responsive neurons, we compared their nuclear transcriptome under different physiological conditions. Analyses of all conditions for our GFP-positive nuclei, found 1,257 genes to be upregulated by leptin and 853 to be downregulated when comparing fasted mice to ones injected every 12 hours (**Fig. 3a**). Gene ontology analyses found enrichment for genes associated with neuronal function (**Supplementary Figure 2b**). As expected, we found that the expression of *Socs3*, a known leptin responsive transcriptional target^24^, was decreased by fasting and rapidly induced by leptin (**Fig. 3b**). Interestingly, the most differentially expressed gene following leptin treatment was ETS variant 6 (*Etv6*) (**Fig. 3b**). Similar to *Socs3, Etv6* is also a known transcriptional target of *Stat3* and is known to inhibit its function in tumor cells^25^, but has yet to be associated with leptin response. Another leptin response gene was guanylate cyclase 2C (*Gucy2c*) (**Fig. 3b**). *Gucy2c* encodes a transmembrane receptor that is expressed in the hypothalamus and causes obesity with an increase in leptin serum levels when deleted in mice^26^. Combined, our results show that transcriptional analysis of FACS sorted nuclei can be used to detect transcriptional changes in small neuronal subpopulations under different physiological conditions.

**Fig. 3.**
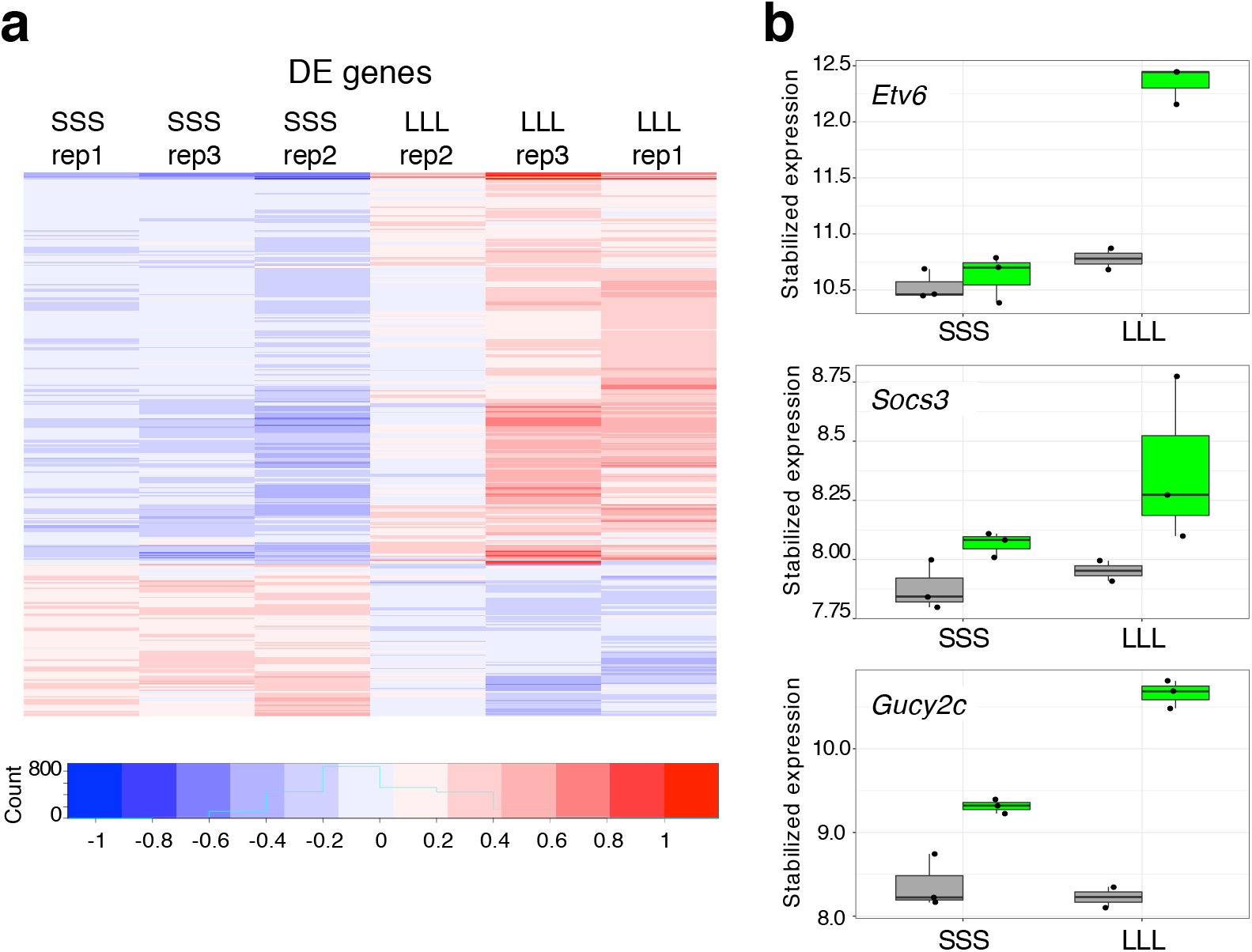
Leptin-responsive transcriptome. **a** A heatmap of mean subtracted, log stabilized read counts showing differentially expressed genes for fasted mice treated either with saline (SSS) or with leptin at 12, 24 and 34 hours (LLL). **b** Boxplots showing the expression levels of *Etv6, Socs3* and *Gucy2c* in LepRb GFP-positive (green) or negative (grey) nuclei within the two conditions. The y-axis shows stabilized expression which is log2 scale for sequencing depth normalized counts determined by DESeq2^39^. Box plots represent 25th to 75th percentiles and the midline indicates the median. n=3 biologically independent replicates.

### Mapping gene regulatory regions in neuronal subpopulations

To identify gene regulatory elements that are active in LepRb expressing neurons, we performed ATAC-seq and ChIP-seq for H3K27ac in sorted nuclei from these neurons. While the starting material for ATAC-seq are nuclei, making the procedure straightforward, ChIP-seq required the development of a novel protocol following nuclei sorting. Briefly, nuclei were first sorted and then isolated via centrifugation in 1M CaCl2 and 1M MgAc2. Gentle fixation using 1% formaldehyde followed by quenching with 125 mM glycine was carried out on the pellet which was subsequently lysed and subjected to ChIP (see Methods for more details).

We identified a total of 32,377 regions that are active in leptin responsive neurons by ATAC-seq and 50,218 regions that are active by H3K27ac ChIP-seq respectively. To confirm the relevance and specificity of the identified elements, we compared our RNA-seq to our ChIP-seq and ATAC-seq results and found that active regulatory regions correlated to levels of gene transcription. Genes near an ATAC-seq or ChIP-seq LepRb enriched peak in leptin responsive neurons were significantly more likely to show the same direction of gene expression changes when compared to peaks with non-significant differences between GFP positive or negative neurons (p-value < 9.3e-7 or 2.2e-16 for ATAC-seq and ChIP-seq respectively; Wilcoxon rank sum test) (**Supplementary Figure 5**). We further examined known leptin pathway associated genes and compared the active regions for these genes in leptin-responsive neurons (from GFP+ nuclei after sorting) to those from all the other hypothalamic cell types (GFP-nuclei after sorting). As specific examples, we observed a strong enrichment of H3K27ac ChIP-seq and ATAC-seq signals specifically in GFP-positive nuclei near the *Lepr* gene promoter, the top positively differentially expressed gene in our RNA-seq compared to all other cell types (**Fig. 2b**) and a reduction near the *Sim1* promoter (**Fig. 4a-b**), one of the top negatively differentially expressed gene (**Fig. 2b**).

**Fig. 4.**
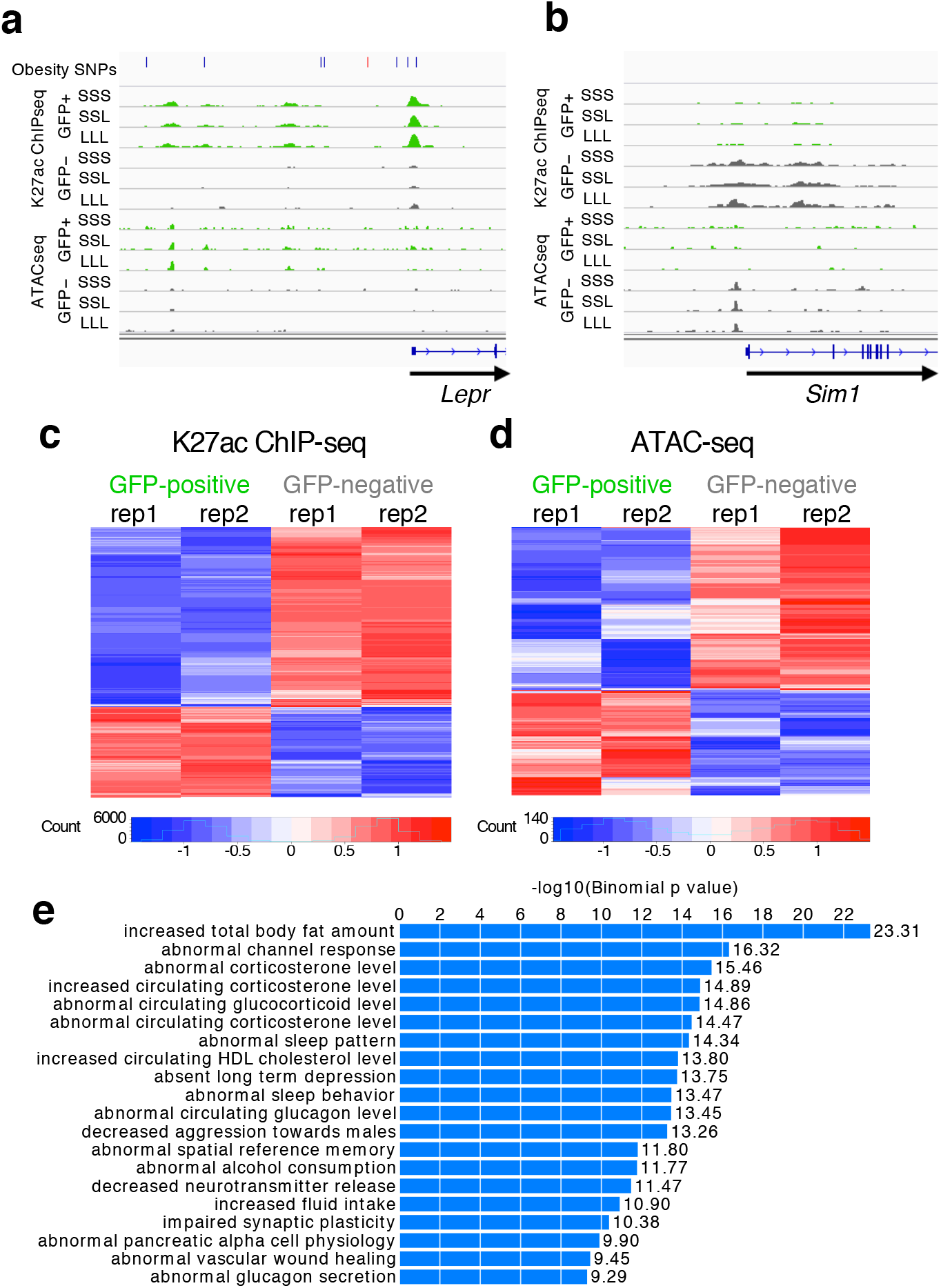
LepRb-GFP positive regulome. **a-b** Genomic snapshots of *Lepr* (**a**) and *Sim1* (**b**) loci showing enrichment of H3K27ac ChIP-seq and ATAC-seq signals around *Lepr* in Leptin-responsive neurons (green) compared to GFP-negative nuclei (grey) and the opposite for *Sim1*. Leptin conditions include mice treated with saline (SSS) or leptin (LLL) injections at 12, 24 and 34 hours (LLL) or saline injections at 12 and 24 hours followed by a single leptin injection at 34 hours (SSL). The obesity SNPs track for *Lepr* shows obesity-associated SNPs^33^, including SNPs that are in linkage disequilibrium (r^2^>0.8) with the lead SNP rs11804091 (red line). Each of the two replicates for ChIP-seq and ATAC-seq presented comparable signals as shown. **c-d** Heatmaps showing differential ChIP-seq (**c**) and ATAC-seq (**d**) peaks between leptin-responsive and GFP negative nuclei. Two replicates are shown for each condition. **e** GREAT analysis^27^ for H3K27ac ChIP-seq peaks that are enriched in LepRb GFP-positive nuclei (n=7,726; p-values obtained by binomial enrichment test).

We then compared peak enrichment between LepRb GFP positive nuclei and all other hypothalamic cell types (GFP negative). We found 17,629 H3K27ac ChIP-seq peaks to be differentially enriched (**Supplementary data file 2**), with 9,903 H3K27ac peaks upregulated in GFP positive nuclei and 7,726 downregulated (**Fig. 4c-d, Supplementary data file 2**). To validate that we are obtaining ChIP-seq peaks that are unique to LepRb GFP positive nuclei, we next used the Genomic Regions Enrichment of Annotations Tool [GREAT^27^]. This tool analyzes the nearby genes of a set of genomic regions and provides biological annotations that are enriched for these genes. Analysis of our H3K27ac upregulated peaks found that the top enriched term for mouse phenotype for the neighboring genes was ‘increased total body fat amount’ (**Fig. 4e**), supporting that we are obtaining a relevant enriched set of genomic peaks.

We next analyzed peaks for transcription factor binding sites (TFBS) enrichment. Using MEME-ChIP^28^ on H3K27ac peaks differentially enriched peaks between GFP-positive and GFP-negative nuclei identified multiple known and novel motifs differentially represented between these two datasets. In the GFP-positive enriched peak set, the STAT4 (MA0518.1) motif was the most overrepresented, with an e-value of 9.1e^−470^ occurring 63,939 times (**Supplementary Table 2**). Another sixteen motifs, including CDX2, NEUROD1, MAX:MYC, SMAD2/3/4, were identified as overrepresented in GFP-positive enriched peaks, but not in GFP-negative enriched peaks. The most highly enriched known motif in the GFP-negative enriched peaks was KLF5 (MA0599.1; e-value 6.7e^−172^; **Supplementary Table 2**). Another seventeen motifs, including CREB3L1, SOX8 and ELK4, were identified as overrepresented in the GFP-negative enriched peaks, but not in the GFP-positive enriched peaks. We also analyzed our data for H3K27ac ChIP-seq or ATAC-seq peaks that are enriched following leptin response [i.e. comparing between chronic leptin injections (LLL) and saline injections (SSS)]. We found 146 H3K27ac ChIP-seq and 28 ATAC-seq peaks to be enriched (raw p-value of 0.01; Wald test) between these two conditions (**Supplementary data file 2**).

### Functional characterization of a *Socs3* enhancer

To further validate whether our LepRb neuron nuclei regulome contains relevant regulatory regions, we set out to test candidate enhancer sequences for their function. We selected three candidate enhancers located downstream of *Socs3* (Socs3-1, Socs3-2, and Socs3-3). These regions were chosen due to their increased enrichment of H3K27ac marks and chromatin accessibility in nuclei of LepRb expressing neurons (**Fig. 5a**). We tested the ability of these enhancers to increase the activity of a minimal promoter driving the expression of luciferase in mHypoA-Pomc cells, an established cell line from mouse hypothalamus Pomc-expressing neurons^29^. From the three sequences, one sequence, Socs3-3, showed significant enhancer activity in this cell type (**Fig. 5b**).

**Fig. 5.**
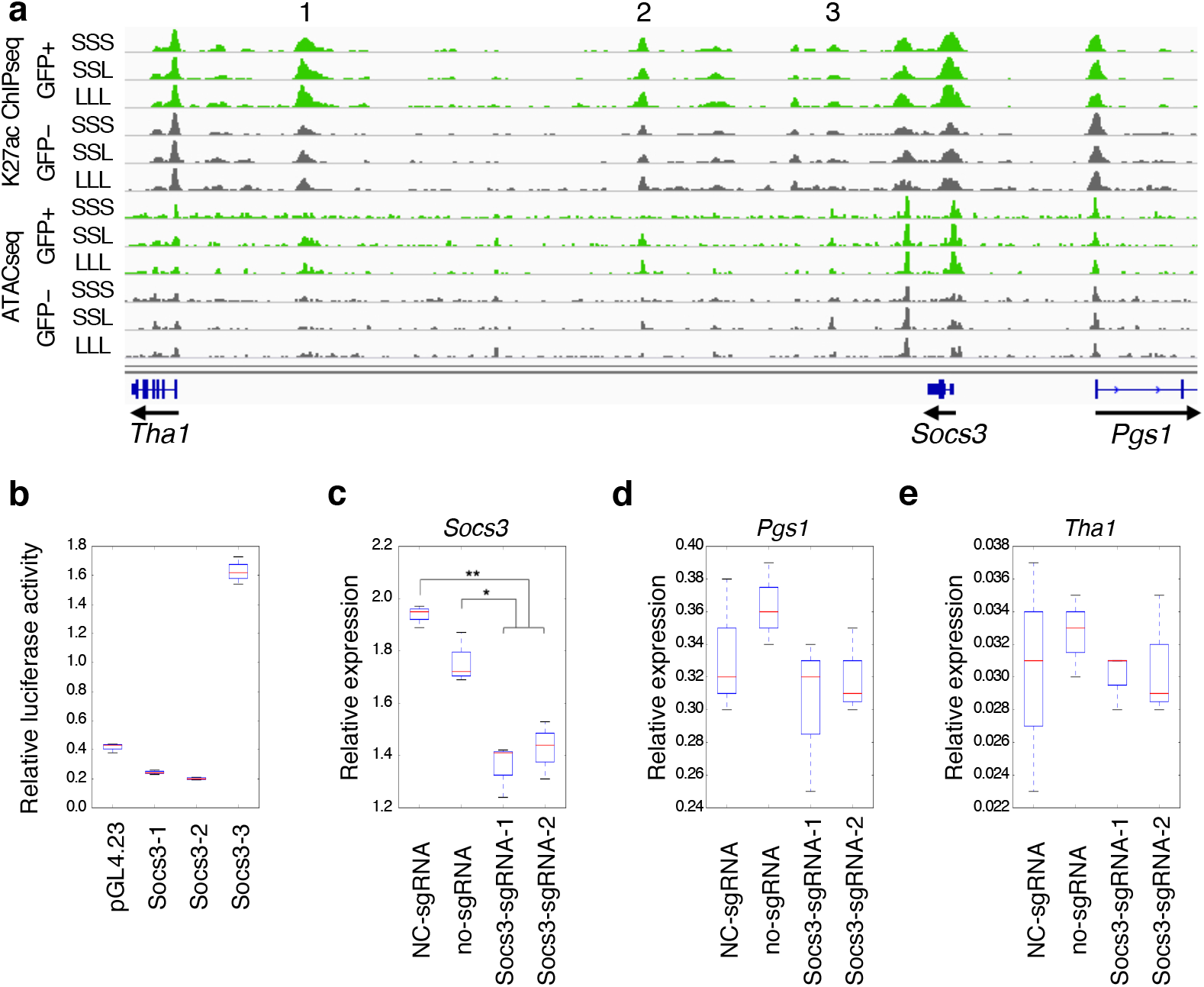
Functional characterization of a *Socs3* enhancer. **a** A genomic snapshot of the *Socs3* locus showing three candidate enhancer sequences, Socs3-1 (1), Socs3-2 (2), and Socs3-3 (3), that are enriched for H3K27ac ChIP-seq and ATAC-seq signals in LepRb GFP-positive nuclei (green) compared to GFP-negative nuclei (grey). Each of the two replicates for ChIP-seq and ATAC-seq presented comparable signals as shown. **b** Luciferase assay for all three candidate enhancer sequences (Socs3-1, Socs3-2, and Socs3-3). Fold activity is compared to the empty vector (pGL4.23) and data are presented as means ± the lower and upper quartile and lines represent the minimum and maximum from three independent biological experiments. **c-e** CRISPRi targeting of Soc3-3. Two sgRNAs (Socs3_sgRNA_1 and Socs3_sgRNA_2) that target the Socs3-3 enhancer along with dCas9-KRAB show significantly reduced expression of *Socs3* via RT-qPCR when compared both to a negative control sgRNA (NC-sgRNA) or no infection control (no-sgRNA). The neighboring genes, *Pgs1* (**d**) and *Tha1* (**e**), do not show any significant changes in gene expression. Boxes represent 25th to 75th percentiles and the midline indicates the median. n=3 biologically independent replicates. Two-sided student’s t-test was used to determine statistical significance (**p<0.01. *p<0.05).

To test whether Socs3-3 regulates *Socs3* expression, we used CRISPR inactivation [CRISPRi^30^] to target this sequence and then measured endogenous *Socs3* expression levels. Using lentivirus, two different sgRNAs targeting the Socs3-3 enhancer were infected into mHypoA-Pomc cells along with a catalytic inactive Cas9 (dCas9) fused to the KRAB repressor. We found that *Socs3* expression was significantly decreased by either sgRNA (**Fig. 5c**). A similar analysis of the neighboring genes around *Socs3*, phosphatidylglycerophosphate synthase 1 (*Pgs1*) and threonine aldolase 1(*Tha1*), did not find any significant change in their expression (**Fig. 5d-e**). Combined, our results confirm that our approach can be used to identify functionally relevant regulatory regions in sub-populations of neurons.

### LepRb neuron regulatory elements overlap obesity-associated SNPs

We next set out to test whether obesity-associated GWAS SNPs could be physically linked to regulatory elements identified in LepRb neurons. We obtained a list of GWAS reported obesity-associated single nucleotide polymorphisms (SNPs) from Ghosh et al^12^ and used Plink^31^ to obtain a list of all SNPs that are in linkage disequilibrium with these tag SNPs using an r^2^ of at least 0.8 as a cutoff (**Supplementary Table 3**). The coordinates of our H3K27ac and ATAC peaks were transferred to the human genome (hg19) using the UCSC LiftOver tool (see Methods). Out of the 61,180 merged peaks, 51,983 (85%) peaks successfully lifted over to hg19. We then tested whether the H3K27ac ChIP-seq or ATAC-seq peaks that were found in LepRb GFP-positive nuclei overlap our list of obesity-associated SNPs. For H3K27ac peaks, we found 196 GFP-positive peaks that overlap obesity-associated SNPs, an observation that was statistically significant (p-value <= 1e^−4^) by a random permutation test carried out by shuffling peak locations 1e^4^ times and excluding ENCODE blacklist regions^32^. Similarly, for ATAC-seq peaks, we found 56 GFP-positive peaks that overlap obesity-associated SNPs. We next compared our H3K27ac peaks to mouse ENCODE peaks after converting them using the UCSC LiftOver tool to human genome (hg19). While many tissue types showed similar enrichment ratios and p-values to the GFP positive dataset, the SNPs and genomic regions of the H3K27ac peaks were distinct across groups (**Figure 6a**). We observed similar results for our ATAC-seq peaks (p-value equal to 3.2e^−3^). Other ATAC-seq datasets^19^ in vasoactive intestinal peptide expressing interneurons (VIP), parvalbumin expressing fast-spiking interneurons (PV) and excitatory pyramidal neurons did not show as significant of an enrichment by this method (p-value equal to 0.012, 0.055, and 0.125, respectively). In summary, our analyses show a unique significant enrichment for obesity-associated SNPs in our predicted regulatory elements.

**Fig. 6.**
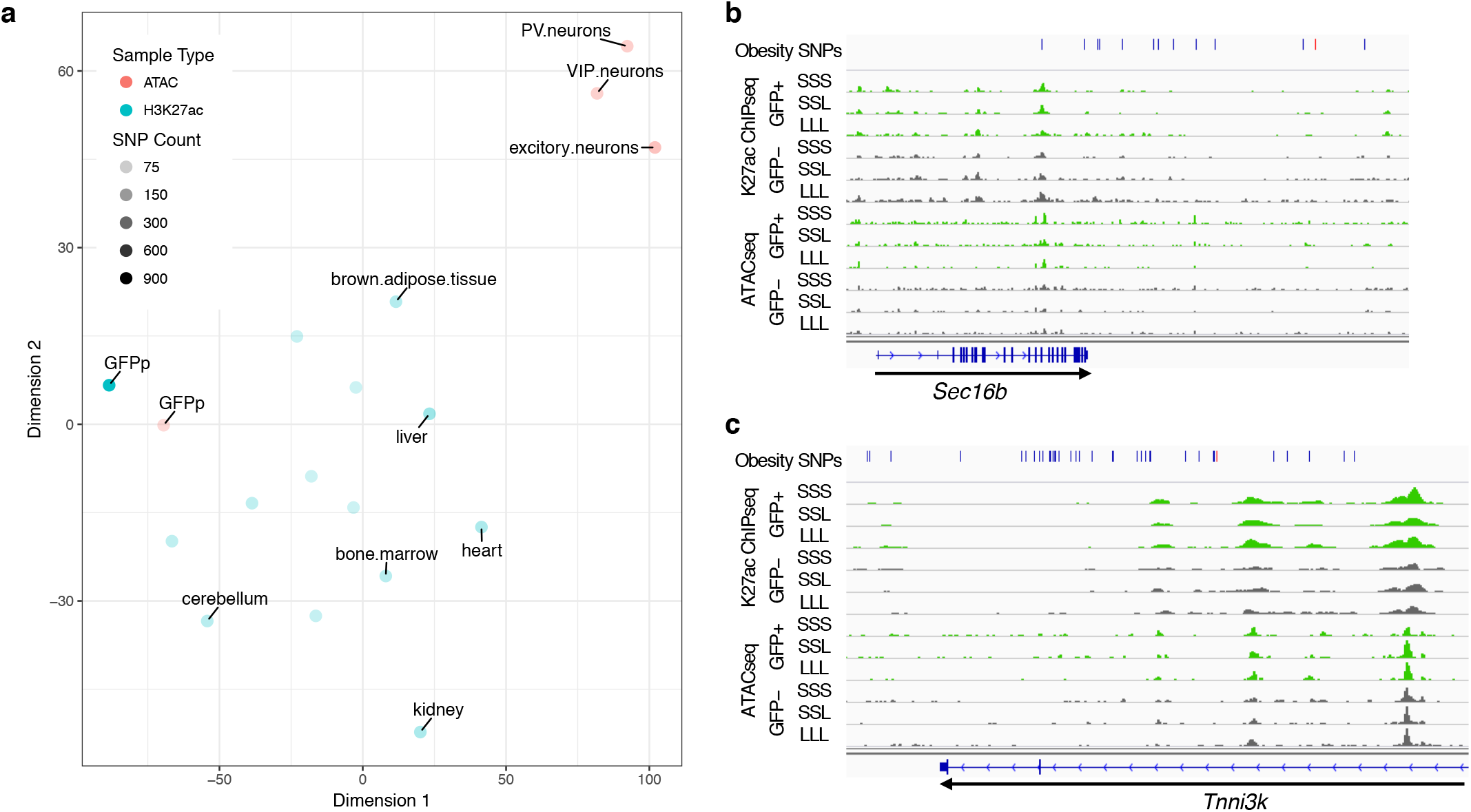
Obesity GWAS loci. **a** tSNE plot showing the differences in obesity-associated SNP-overlaps between tissue and data types. The tSNE plot was generated from a matrix of obesity-associated SNPs which overlap each ChIP-seq or ATAC-seq dataset, such that rows were SNPs and columns samples. Each field was assigned either a 0 (no overlap) or 1 (obesity-associated SNP was overlapped). Color represents the data type: Red – ATAC-seq, Blue – ChIP-seq. The darkness of the color represents the total number of obesity-associated SNPs that overlapped each dataset. **b-c** Genomic snapshots of *Sec16b* (**b**) and *Tnni3k* (**c**) loci showing H3K27ac ChIP-seq and ATAC-seq signals in leptin-responsive neurons (green) compared to GFP-negative nuclei (grey). Leptin conditions include mice treated with saline (SSS) or leptin (LLL) injections at 12, 24 and 34 hours (LLL) or saline injections at 12 and 24 hours followed by a single leptin injection at 34 hours (SSL). The obesity SNPs track shows the obesity-associated lead SNP a as red line and in addition SNPs that are in linkage disequilibrium (r^2^>0.8) as black lines. Each of the two replicates for ChIP-seq and ATAC-seq presented comparable signals as shown.

We also examined a number of specific GWAS loci to determine whether the Leprb regulome could help identify obesity-associated functional SNPs. For example, the *Lepr* locus (**Fig. 4a**), where SNPs were found to be associated with obesity^33^, showed an overlap between these SNPs and LepRb regulatory elements. Other examples include the SEC16 homolog B, endoplasmic reticulum export factor (*SEC16B*), SH2B adaptor protein 1 (*SH2B1*), gastric inhibitory polypeptide receptor (*GIPR*) and TNNI3 interacting kinase (*TNNI3K*) loci (**Fig. 6b-c; Supplementary Figure 6**), all of which have been highly associated with obesity through GWAS^11^.

We also tested whether our LepRb neurons overlap hypothalamus expression quantitative loci (eQTLs) as determined by the Genotype-Tissue Expression (GTEx) project^34^. We found a significant enrichment (<1e^−3^ p-value; random permutation test for 1000 data shuffles excluding ENCODE black list regions^32^) for eQTLs in both our H3K27ac ChIP-seq and ATAC-seq peaks (**Supplementary data file 3**). These results show that our characterized LepRb neuron nuclei regulatory elements can be used as a blueprint to identify putative functional obesity-associated variants.

## Discussion

Numerous GWASs have been performed to find common genetic variants associated with obesity, and over 250 loci have been identified in which clusters of linked SNPs predispose to an increase in BMI^10^. However, little progress has been made in outlining the causal SNPs and understanding the mechanisms by which they affect the phenotype. Most of these SNPs are found in noncoding regions of the genome and maps of relevant functional elements potentially affected by these SNPs are required. In particular, since obesity-associated loci appear to be most often located near genes expressed in the central nervous system, the capacity to map genomic regulatory regions in subpopulations of neurons implicated in the control of energy homeostasis would be a first step towards linking obesity associated SNPs with their function.

Here, using LepRb expressing neurons as an example, we developed a unique protocol that can identify gene regulatory elements from a relevant sub-population of neurons thus providing a resource to start addressing the functional consequences of human obesity-associated SNPs. First, we demonstrate that FACS sorting of a genetically labeled sub-population of hypothalamic neurons results in a highly enriched nuclei population. This method is dependent on the expression of a nuclei anchored GFP in the relevant population of neurons and could therefore be used for any neuronal population for which a Cre reporter is available.

The characterized LepRb expressing neuron population is still heterogeneous and is from both genders (does not account for sex-specific differences). More specific sub-populations of Cre expressing neurons could be isolated by nuclei labeling through stereotaxic injection of a Cre dependent (DIO) Adeno Associated Virus (AAV) expressing SUN1 sfGFP. For example, this method could be used to find regulatory regions in LepRb expressing neurons that are more specific to the arcuate nucleus or the dorsomedial hypothalamic nucleus. This approach also provides the opportunity of labelling the nuclei that are expressing Cre only at the time of injection. Finally, if a Cre reporter is not available, stereotaxic injection of AAVs expressing Cre recombinase under the control of a synapsin promoter, that confers gene expression in neurons, can also be used to label anatomical subsets of neuronal nucleis in *R26-CAG-LSL-Sun1-sfGFP-myc* mice. We also developed a unique protocol that provides the ability to carry out ChIP-seq for H3K27ac on these sorted nuclei. We demonstrate the fidelity of this map to transcriptional activity and its functional relevance. We did not however identify many leptin treatment enriched peaks. This is likely due to technical and statistical constraints, using an extremely low number of nuclei in these assays (**Supplementary Table 1**) leading to our inability to generate statistically significant comparisons. Further development of this protocol could allow increased sensitivity and generalization to the study of additional histone marks or more specific transcription factors.

Demonstrating that our map will be useful for linking genotype with an obesity phenotype, we highlight the overlap between our LepRb neurons H3K27ac peaks and GWAS SNPs at a subset of obesity-associated loci, including some with strong BMI associations. Our characterized regulatory elements provide candidate sequences for further functional analyses of these obesity-associated loci. Of note and as expected, several obesity GWAS peaks did not overlap our LepRb neuron regulatory elements, underlying the necessity to establish regulatory maps of other hypothalamic neuronal populations implicated in body weight regulation such as MC4R expressing neurons. The two-pronged approach we describe to demonstrate the functional relevance of identified SOCS3 enhancers can also be further adapted to the study of the functional effects of candidate obesity-associated SNPs on regulatory function. Finally, we also provide evidence that nuclear transcriptomes that can be obtained following nuclei sorting are highly specific, can be used to study the effect of physiological changes on transcriptional activity in specific neuronal populations, and unlike TRAP-seq allow for the study of untranslated RNA.

## Acknowledgments

We thank Dr. Jessica Tollkuhn (CSHL) for critical advice and Dr. Martin Myers (U Michigan) for sharing LepRb TRAP-seq data. We also thank Dr. Aaron Hardin and Sawitree Rattanasopha for help with dissections and Drs. Marielle Cavrois and Mekhala Maiti for assistance with FACS. This article was supported in part by a grant from the National Institute of Diabetes and Digestive and Kidney Diseases (NIDDK) 1R01DK090382 (for N.A. and C.V) and the UCSF Nutrition Obesity Research Center funded by that National Institute of Health P30DK098722. N.A. is also supported by grants by the National Human Genome Research Institute (NHGRI) and Division of Cancer Prevention, National Cancer Institute grant number 1R01CA197139, National Institute of Mental Health grant number 1R01MH109907, National Institute of Child & Human Development 1P01HD084387, NHGRI grant number 1UM1HG009408 and National Heart Lung and Blood Institute 1R01HL138424. The Gladstone Institute flow cytometry core is supported by NIH P30 AI027763 for using LSR2, Calibur, VYB, Aria, HTFC and ImageStream, NIH S10 RR028962 and the James B. Pendleton Charitable Trust for using the FACSAria cell sorter, and DOD W81XWH-11-1-0562 for using an ImageStream.

## Author Contributions

F.I. C.V. and N.A. conceived and designed the study. F.I. performed the mouse, genomic and enhancer experiments. K.K.M. and N.M. helped with mouse dissections. Y.W. carried out the immunostaining and W.L.E. preformed the computational analyses. F.I. W.L.E. C.V. and N.A. analyzed the data, C.V. and N.A. provided resources and critical suggestions and F.I. W.L.E. C.V. and N.A. wrote the manuscript.

## Methods

### Mice

LepRb^cre^ (Lepr^tm3(cre)Mgmj^)^18^ heterozygote mice were crossed to SUN1-sfGFP (Gt(ROSA)26Sor^tm5(CAG-Sun1/sfGFP)Nat^)^19^ homozygote mice, both on a mixed 129S1/Sv and C57BL/6J background, to generate LepR^cre^/+; SUN1-sfGFP/+ mice. Mice were genotyped by PCR using primer sequences described in **Supplementary Table 4**. Mice were fed *ad libitum* Picolab mouse diet 20, 5058 containing 20% protein, 9% fat, 4% fiber for whole study. Calories provided by: protein 23.210%, fat (ether extract) 21.559% and carbohydrates 55.231%. LepR^cre^/+; SUN1-sfGFP/+ mice between 2-4 months old were then randomly chosen (see **Supplementary Table 1** for number of mice and genders used) for the following treatments: 1) Fed *ad libitum* for 36 hours; 2) Fasted for 36 hours and injected with saline three times (12, 24, and 34 hours after the beginning of fasting); 3) Fasted for 36 hours and injected with leptin (5 mg/kg body weight) three times (12, 24, and 34 hours after the beginning of fasting); 4) Fasted for 36 hours and injected with leptin (5 mg/kg body weight) (Los Angeles Biomedical Research Institute) at 34 hours. The mice were euthanized and the hypothalami were dissected. All hypothalami used in the following experiments were confirmed for GFP expression under a fluorescent microscope. All animal work was approved by the UCSF Institutional Animal Care and Use Committee.

### Immunostaining

Mice were anesthetized with pentobarbital (7.5 mg/0.15 ml, i.p.) and transcardially perfused with 10ml of heparinized saline (10 U/ml, 2 ml/min) followed by 10ml of phosphate-buffered 4% paraformaldehyde (PFA). Brains were removed, postfixed for 24 hours in 4% PFA, and then equilibrated in PBS with 30% sucrose for 72 hours. Brains were sectioned coronally (35 microns) on a sliding microtome (Leica SM 2000R). Immunohistochemistry was performed as previously described ^35–37^. Coronal brain sections that had been stored in PBS at 4°C were permeabilized and blocked with 3% normal goat serum/0.3% Triton X-100 for 1 hour and then incubated at 4°C overnight with chicken anti-GFP (abcam, ab13970) as the primary antibody and goat anti chicken IgY Alexa fluor 488 (Invitrogen, A11039) as the secondary antibody at a dilution of 1:500. Sections were placed in 4,6-diamidino-2-phenylindole (DAPI) (0.2 g /ml; 236276; Roche) for 10 minutes and then mounted on plus coated slides and coverslipped using Vectashield (H-1000; Vector Laboratories). Images were captured using a Zeiss Apotome.

### RNA-seq

Three biological replicates were done for each condition and five hypothalami were used in each replicate. Hypothalami were pooled and homogenized by douncing 30 times in 1 mL lysis buffer (15 mM Tris-HCL (pH 7.5), 0.1 mM EGTA, 15 mM NaCl, 60 mM KCl, 5 mM MgCl2, 300 mM sucrose, 1 mM DTT, 0.1 mM PMSF, 0.15 mM spermine, 0.5 mM spermidine, Complete protease inhibitor (Roche) and 20mM Na-butyrate (Sigma), RiboLock RNase inhibitor) using a tissue grinder type B (Kimble Chase). The cell lysate was centrifuged at 500 g for 5 minutes at 4°C. Pellet was resuspended in 1 mL lysis buffer, and equal volume of nuclei isolation buffer (lysis buffer supplemented with NP40 at 0.4%) was added. The lysate was incubated on ice for 5 minutes, followed by centrifugation at 7000 g for 5 minutes at 4°C. Cell nuclei pellet was re-suspended in 250 uL FACS buffer (0.15 mM spermine, 0.5 mM spermidine, Complete protease inhibitor (Roche) and 20mM Na-butyrate (Sigma), RiboLock RNase inhibitor in PBS. FACS was performed with FACS Aria II (BD Biosciences) equipped with 70 um nozzle. 50,000-70,000 GFP positive and negative nuclei were sorted directly into buffer RLT supplemented with beta-mercaptoethanol. Cell nuclei with fluorescence levels 0-600 were defined as GFP negative, and those with ≥6,000 as GFP positive (**Supplementary Figure 1**). RNA was extracted using RNeasy mini kit (Qiagen) following the manufacture’s protocol. RNA was quantified with Qubit RNA HS assay kit. cDNA was amplified using Ovation V2 kit (NuGEN) and sequencing libraries were generated using NexteraXT kit (Illumina). RNA-seq was carried out on an Illumina HiSeq 4000. Sequence alignment was performed using STAR^38^. Mappings were restricted to those that were uniquely assigned to the mouse genome and unique read alignments were used to quantify expression and aggregated on a per-gene basis using the Ensembl (GRCm37.67) annotation. We analyzed these raw data using DESeq2^39^ to assess variance and differential expression between sample groups. All RNA-seq data was deposited in NCBI as Bioproject PRJNA418102. To compare our RNA-seq result to previous work isolating LepRb neurons followed by TRAP-seq^22^, we selected all differentially expressed genes between GFP positive and GFP negative conditions. We then further divided those genes into genes more highly expressed in GFP positive cells or higher in GFP negative cells. These differentially expressed genes where then compared to the TRAP-seq results in LepRb positive or LepRb negative cells. Additionally, we carried out gene ontology and pathway enrichment analysis on differentially expressed genes using DAVID^40^ through the DAVIDWebService package^41^ on Bioconductor. For unsupervised clustering, we utilized the WGCNA^20^ package. Genes were first filtered to remove those with less than 90% of the samples containing normalized read count depth of 20 (kept 11,125 genes). Module assignment was then inspected after carrying out the blockwiseModules function with a parameter sweep for options power (values: 6, 7, 8 and 9), minimumModuleSize (values 10, 15, 20, 25, 30, 40 and 50) and mergeCutHeight (values .1, .15, .20, .25, .30 and .35). From this power = 7, minModuleSize = 50 and k = .15 were determined to give optimal module assignment based on inspection of gene dendrograms, the number of total clusters and the number of genes in each cluster. From this point, the modules were correlated to the experimental groups following the WGCNA vignette and p-values of this correlation were adjusted for multiple testing using the False Discovery Rate (FDR). As with the differentially expressed genes, enrichment for gene ontology and pathways in each module’s gene set was determined using DAVID^40^ and the DAVID WebService package^41^.

### ChIP-seq

Twenty hypothalami were used for each individual ChIP-seq experiment and two biological replicates were done for each condition. Cell nuclei were isolated and pelleted as described above. Cell nuclei pellet was re-suspended in 2 mL FACS buffer (0.15 mM spermine, 0.5 mM spermidine, Complete protease inhibitor (Roche) and 20mM Na-butyrate (Sigma) in PBS). FACS was performed with FACS Aria II (BD Biosciences) equipped with 100 um nozzle. 300,000-540,000 GFP positive and negative nuclei were sorted into PBS. After FACS, FACS buffer was added for a final volume of 10 mL. 50 uL of 1M CaCl2 and 30 uL of 1M MgAc2 was added and incubated on ice for 5 minutes, followed by centrifugation at 1800 g for 15 minutes at 4°C. The supernatant was removed and FACS buffer supplemented with 1% formaldehyde was gently added onto the pellet to crosslink chromatin. The crosslinked pellet was then washed with FACS buffer supplemented with 125 mM glycine without disturbing the pellet. The crosslinked chromatin was lysed in 130 uL of Buffer B (LowCell# ChIP kit, diagenode) supplemented with Complete protease inhibitor (Roche) and 20mM Na-butyrate (Sigma). The lysed chromatin was sheared using a Covaris S2 sonicator to obtain on average 250bp size fragments. 870 uL of Buffer A (LowCell# ChIP kit, diagenode) supplemented with complete protease inhibitor (Roche) and 20mM Na-butyrate (Sigma) was added. 20 uL of the chromatin solution was saved as an input control. A mixture of 40 uL of Dynabeads protein A and 40 uL of Dynabeads protein G was washed twice with Buffer A (LowCell# ChIP kit, diagenode) and resuspended in 800 uL of Buffer A. 10 ug of H3K27ac antibody (ab4729, Abcam) was added to the beads, and gently agitated at 4°C for 2 hours. The beads-antibody complex was precipitated with a magnet, and the supernatant was removed. 800 uL of shared chromatin was added to the beads-antibody complex and rotated at 4°C overnight. The immobilized chromatin was then washed with Buffer A three times and Buffer C once, and eluted in 100 uL of IPure elution buffer (IPure kit, diagenode). In addition, 80 uL of IPure elution buffer was added to 20 uL input. DNA was purified using the IPure kit according to manufacture’s protocol. Sequencing libraries were generated using Accel-NGS 2S Plus DNA library kit (Swift Biosciences). DNA was quantified with Qubit DNA HS assay kit and Bioanalyzer (Agilent) using the DNA High Sensitivity kit. Massively parallel sequencing was performed on an Illumina Hiseq 4000. Sequencing reads were mapped to the genome using bowtie^42^ allowing one mismatch per read-alignment and only uniquely aligned reads (-v 1 -m 1). Peaks were called against input using MACS2^43^. Reliable peaks were identified using the ENCODE Irreproducible Discovery Rate (IDR; ^44^) pipeline, which establishes a p-value cutoff to accept peak calls for each condition. For differential peak intensity analysis, peaks across conditions were partitioned using BEDTools^45^ and read coverage obtained using HTSeq^46^. Peaks differentially enriched for H3K27ac histone marks were then identified using DESeq2^39^. Differentially enriched peaks were tested for enrichment for novel motifs and known transcription factor binding sites with MEME-ChIP^28^. Test peak regions were split into peaks either showing increased enrichment in GFP-positive cells or GFP-negative cells. The peaks were then divided into 500bp windows or smaller using BedTools^45^ makewindows command. The Fasta sequences for these regions were then extracted using BedTools getfasta with the repeat masked version of the Ensembl Mouse genome. MEME-ChIP was then run on each set of windowed peaks using the following parameters: -db ~/tools/meme-5.0.0/db/motif_databases/JASPAR/JASPAR2018_CORE_vertebrates_non-redundant.meme -meme-minw 6 -meme-maxw 12 -meme-nmotifs 500 -meme-minsites 50 -meme-maxsites 500 -meme-p 12. All ChIP-seq data was deposited in NCBI as Bioproject PRJNA418098.

### ATAC-seq

Five hypothalami were used for the ATAC-seq using two biological replicates for each condition. Cell nuclei were isolated and pelleted as described above. Cell nuclei pellet was re-suspended in 250 uL FACS buffer (0.15 mM spermine, 0.5 mM spermidine in PBS). FACS was performed with the FACS Aria II (BD Biosciences) equipped with 70 um nozzle. GFP positive and negative nuclei were sorted into PBS. After FACS, 49 uL of sorted nuclei (11,200-18,000 nuclei) were mixed with 50 uL Tagment DNA buffer (Nextera DNA sample preparation kit, Illumina) and 1 uL of Tagment DNA enzyme (Nextera DNA sample preparation kit, Illumina), followed by incubation at 37°C for 30 minutes. Tagmented DNA was purified with MinElute reaction cleanup kit (Qiagen). The DNA was size-selected using SPRIselect (Beckman Coulter) according to manufacture’s double size selection protocol. DNA:SPRIselect ratio was 5:3 for right side, and 2:3 for left side selection. Library amplification was performed as described previously^47^. Amplified library was further purified with SPRIselect as described above. DNA was quantified using the Qubit DNA HS assay kit and Bioanalyzer (Agilent) using the DNA High Sensitivity kit. Massively parallel sequencing was performed on an Illumina Hiseq 4000. Sequencing reads were mapped to the genome using Bowtie2^48^; options: --no-unal -X 2000 --no-discordant --no-mixed --local --very-sensitive-local). After duplicate read removal, samples were merged into a GFP positive and GFP negative pool for peak calling with MACS2^43^ and reliable peaks were identified using the ENCODE IDR^44^ pipeline. As with ChIP-seq, peaks were then partitioned with BEDTools^45^, read coverage obtained using HTSeq^46^ and differentially enriched peaks identified using DESeq2^39^. Differentially enriched ATAC-seq peaks were tested for enrichment for novel motifs and known transcription factor binding sites with MEME-ChIP^28^ in a similar fashion as the ChIP-seq peaks above. However, due to peak regions being generally smaller, peaks were divided into 200bp windows or smaller using BedTools^45^ makewindows command. All ATAC-seq data was deposited in NCBI as Bioproject PRJNA418098.

### GWAS and GTEX analyses

Obesity-associated variants were obtained from Ghosh and Bouchard, Nature Reviews Genetics, 2017^12^. Variants in high linkage disequilibrium (r^2^ of at least 0.8) with obesity-associated SNPs were then obtained the 1000 Genomes data^49^ in all five available populations (AMR, AFR, EUR, EAS, SAS) with Plink^31^. All lead and linkage variants from all five populations were then concatenated into a single list with their locations in the human hg19 assembly. The coordinates of these variants were then intersected with the ChIP-seq and ATAC-seq peaks after using the UCSC LiftOver tool using BEDTools^45^. We used a random permutation test to determine the statistical significance of the rate of intersection. To do this, we randomly shuffled lifted over mouse peaks using BEDTools^45^, after excluding regions on the ENCODE^32^ blacklist, a total of 10,000 times. For each shuffle, we recorded the number of peaks that contained an obesity-associated lead or linked variants. We then compared the observed occurrences in the ChIP-seq or ATAC-seq data to the random permutations. The raw p-value was determined to be equal to the number of permutations greater than the observed plus one, divided by the total number of permutations plus one. Raw p-values were adjusted for multiple testing using FDR methods and the R p.adjust function. Hypothalamus eQTL variants were downloaded from the GTEX^50^ portal (https://gtexportal.org/home/datasets). Similar to the obesity-associated variants identified through GWAS studies, we intersected these variants on the human genome with the lifted over ChIP-seq and ATAC-seq peak regions and carried out a random permutation test with a total of 1,000 permutations (fewer permutations were carried out due to the lower degree of significance identified). Raw p-values and FDR adjusted p-values were obtained as described above.

### Luciferase assays

The three genomic regions around *Socs3* (Socs3-1, Socs3-2, and Socs3-3) were amplified via PCR from the mouse genome using specific primers (**Supplementary Table 4**), and cloned into the pGL4.23 (Promega) enhancer assay vector using In-Fusion HD cloning kit (Takara) following the manufacturer’s protocol. For luciferase assays, mHypoA-POMC/GFP-1 cells were seeded in 96-well plate with 1×10^4^ cells/well at 24 hours before transfection. 120 ng of plasmid per well were transfected along with 30 ng of pGL4.74 Renilla luciferase expression vector (Promega), to correct for transfection efficiency, using XtremeGENE HP DNA transfection reagent (Roche). DNA:XtremeGENE ratio was 1:2. Firefly and Renilla luciferase activities were measured 24 hours after transfection using the Dual-luciferase reporter assay system (Promega) on a Synergy2 (Biotek) following the manufacturer’s protocol.

### CRISPRi

sgRNA sequences (**Supplementary Table 4**) were cloned into the pLG1 plasmid (gift from Prof. Jonathan Weissman). Intact pLG1 plasmid that contains sgRNA sequence against EGFP gene was also used as a negative control. sgRNA containing pLG1 vectors along with the pHR-SFFV-KRAB-dCas9-P2A-mCherry (Addgene, plasmid #60954) were used to generate lentivirus using the Lenti-Pac HIV expression packaging kit (GeneCopoeia). Lentivirus was concentrated using the Lenti-X Concentrator (Takara). mHypoA-POMC/GFP-1 cells were infected with sgRNA and KRAB-dCas9 virus (1:1 ratio) at a multiplicity of infection (MOI) of 0.5. Following 48 hours, RNA was extracted using the RNeasy mini kit (Qiagen) and reverse-transcribed using Superscript III reverse transcriptase (Invitrogen). Gene expression was examined by RT-qPCR using SSO fast EvaGreen supermix (Bio-Rad) was carried out on an Eppendorf Realplex 2. Primers used for the qPCR are shown in **Supplementary Table 4**.

### Reporting Summary

Further information on research design is available in the Nature Research Reporting Summary linked to this article.

### Data availability

RNA-seq data are available in Bioproject: PRJNA418102. ChIP-seq and ATAC-seq are available in Bioproject: PRJNA418098. The data that support the findings of this study are available from the corresponding author upon request.

