## Supplementary material for "Genomic and epigenomic mapping of leptin-responsive neuronal populations involved in body weight regulation"

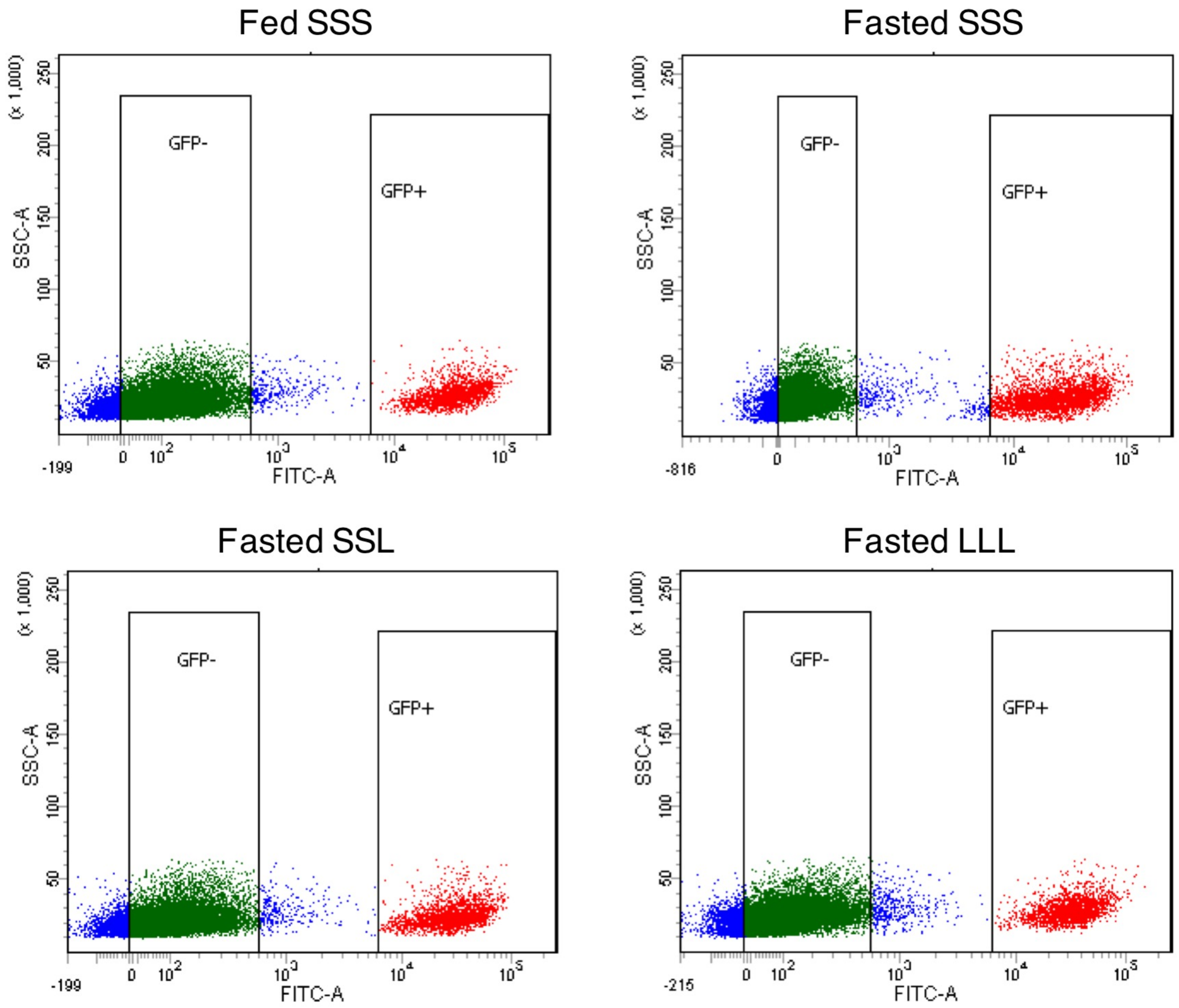


**Supplementary Figure 1. Gating strategy for flow cytometry. a-d** Cell nuclei obtained from mice with four leptin conditions including fed and injected with saline three times (Fed SSS; **a**) fasted and injected with saline three times (Fasted SSS; **b**), saline at 12 and 24 hours and leptin at 34 hours (Fasted SSL; **c**), and fasted and injected with leptin three times at 12, 24 and 34 hours (Fasted LLL; **d**). Cell nuclei with fluorescence activity between zero to 600 were defined as GFP negative, and those with more than 6,000 as GFP positive.

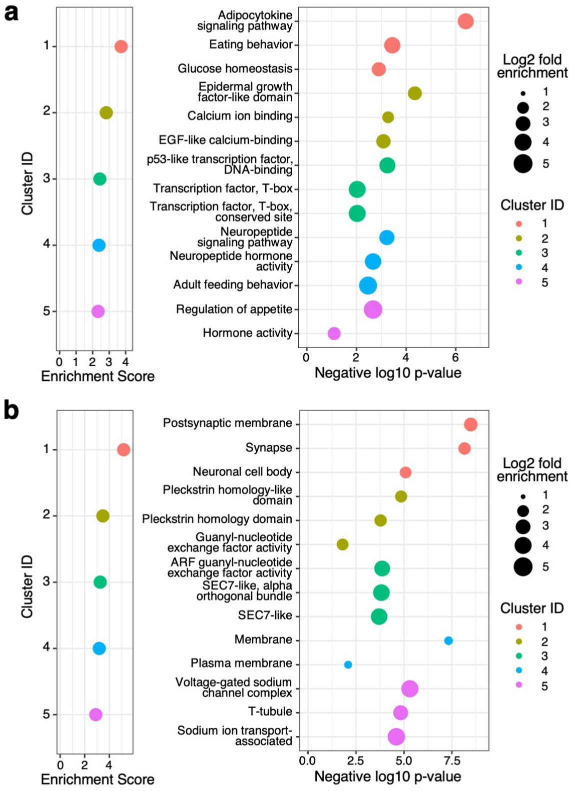


**Supplementary Figure 2. Gene ontology analysis. a** Gene ontology analysis of differentially expressed genes between LepRb GFP positive versus GFP negative nuclei using functional annotation clustering in DAVID[1]. The panel on the left shows the enriched gene ontology clusters and the right panel the terms associated with these clusters. **b** Functional annotation clustering using DAVID[1] of LepRb GFP positive nuclei of genes that are differentially expressed between fasted mice injected at 12, 24 and 34 hours with saline (SSS) or leptin (LLL). The panel on the left shows the enriched gene ontology clusters and the right panel the terms associated with these clusters. N=3 biologically independent replicates, p-values by modified Fishers Exact test (**a, b**).

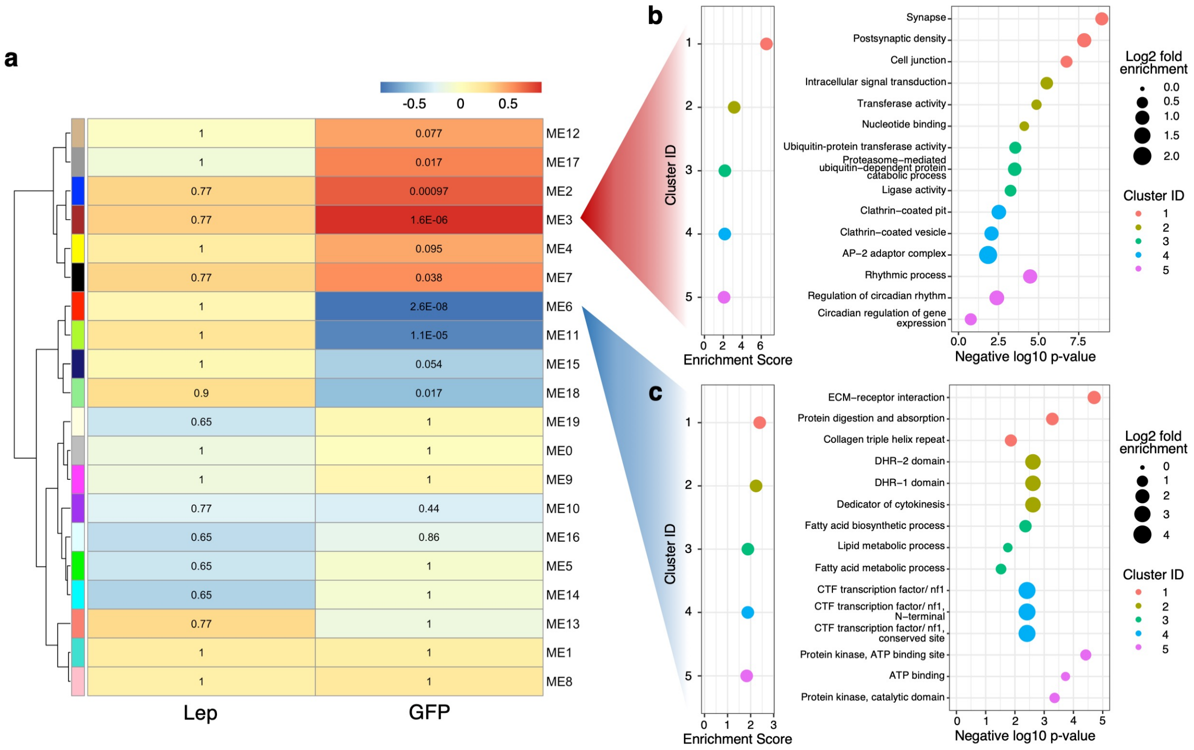


**Supplementary Figure 3. WGCNA analysis. a** WGCNA analysis found 10 modules to be highly correlated with GFP-positive and GFP-negative cell states. **b-c** Gene ontology analysis using DAVID[1] for two of the most significantly correlated modules, ME3 (**b**) and ME6 (**c**). N=3 biologically independent replicates, p-values calculated by Student's asymptotic test and corrected for multiple testing using Benjamini & Hochberg method (**b, c**).

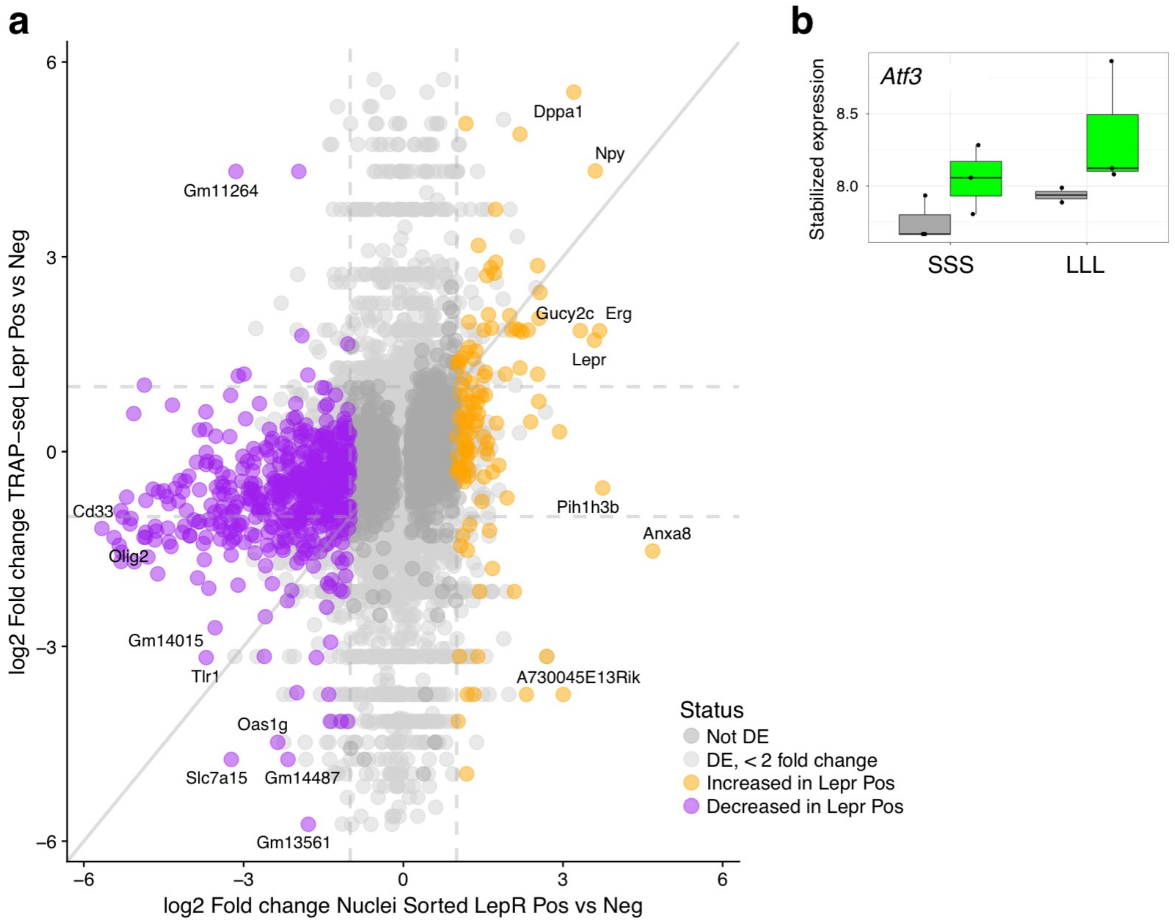


**Supplementary Figure 4. MM plot of TRAP-seq versus our RNA-seq and Atf3 expression levels. a** Comparison of the log2 fold changes between LepRb GFP positive and negative sorted nuclei in this data set (x-axis) to the log2 fold change between LepRb positive verus negative cells in the TRAP-seq assay[2] (y-axis). **b** Boxplot showing the expression level of *Atf3* in LepRb GFP-positive (green) or negative (grey) nuclei within the two conditions. The y-axis shows stabilized expression which is log2 scale for sequencing depth normalized counts determined by DESeq2[3]. Box represents 25th to 75th percentiles and the middle line indicates the median. TRAP-seq N=1 per group, Nuclei Sorted RNA-seq N=3 per group, P-values given by Wald test and corrected for multiple testing by the Benjamini & Hochberg procedure.

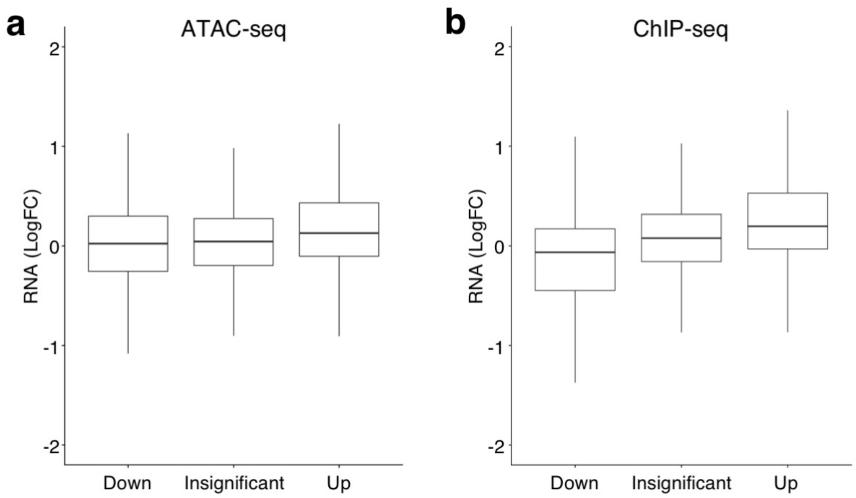


**Supplementary Figure 5. ATAC-seq or ChIP-seq to RNA-seq comparisons. a-b** Comparison between differentially enriched ATAC-seq (**a**) or ChIP-seq (**b**) peaks of leptin-responsive neurons that were upregulated, insignificant or downregulated and nearest gene fold change. Boxes represent 25th to 75th percentiles and the midline indicates the median. ATAC-seq down (N=3745), Insignificant (N=48330), Up (N=3767). ChIP-seq down (N=9903), Insignificante (N=121669), Up (N=7726).

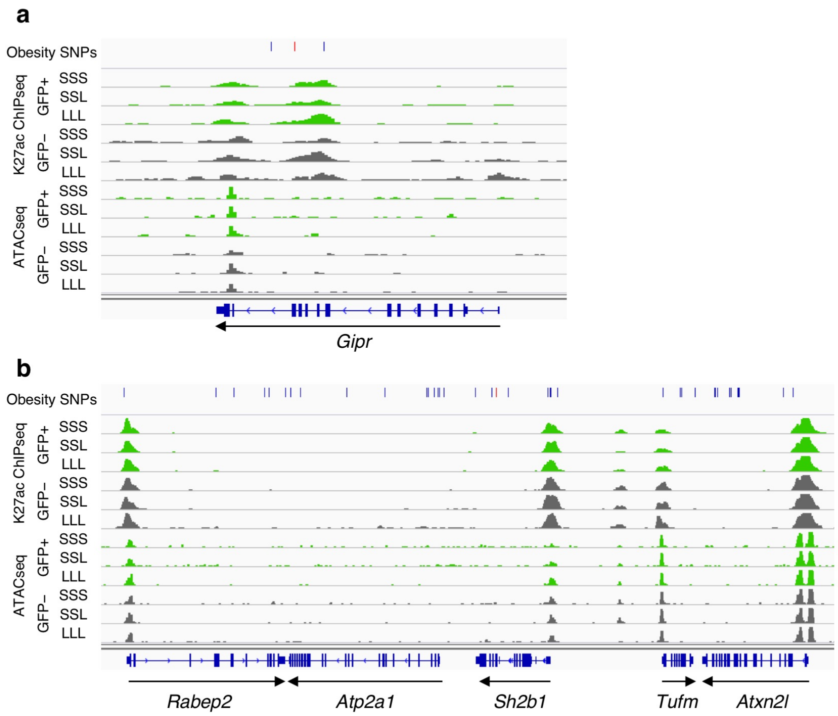


**Supplementary Figure 6. Obesity GWAS loci. a-b** Genomic snapshots of *Gipr* (**a**) and *Sh2b1* (**b**) loci showing H3K27ac ChIP-seq and ATAC-seq signals in leptin-responsive neurons (green) compared to GFP-negative nuclei (grey). Leptin conditions include mice treated with saline (SSS) or leptin (LLL) injections at 12, 24 and 34 hours (LLL) or saline injections at 12 and 24 hours followed by a single leptin injection at 34 hours (SSL). The obesity SNPs track shows the obesity-associated lead SNPs as a red line and in addition SNPs that are in linkage disequilibrium ($r^2$>0.8) as black lines. Each of two replicates for ChIP-seq and ATAC-seq presented comparable signals as shown (**a**, **b**).

### Supplementary Table 1. Number of mice and nuclei used for experiments.

| | | | hnRNA-seq | | ATAC-seq | | H327ac ChIP-seq | |
|---|---|---|---|---|---|---|---|---|
| | | | # mice used (Male, Female) | # nuclei used (thousands) | # mice used (Male, Female) | # nuclei used (thousands) | # mice used (Male, Female) | # nuclei used (thousands) |
| GFP positive | Fed Saline x3 | rep1 | (3, 2) | 50 | (3, 2) | 11.2 | (10, 10) | 568 |
| | | rep2 | (3, 2) | 58 | (2, 3) | 14.4 | (10, 10) | 536 |
| | | rep3 | (3, 2) | 44 | | | | |
| | Fas Saline x3 | rep1 | (3, 2) | 50 | (3, 2) | 11.2 | (10, 10) | 497 |
| | | rep2 | (3, 2) | 56 | (2, 3) | 14.4 | (10, 10) | 540 |
| | | rep3 | (3, 2) | 74 | | | | |
| | Fas Leptin x1 | rep1 | (3, 2) | 50 | (3, 2) | 11.2 | (10, 10) | 435 |
| | | rep2 | (3, 2) | 55 | (2, 3) | 14.4 | (10, 10) | 454 |
| | | rep3 | (3, 2) | 61 | | | | |
| | Fas Leptin x3 | rep1 | (3, 2) | 50 | (3, 2) | 11.2 | (10, 10) | 477 |
| | | rep2 | (3, 2) | 50 | (2, 3) | 18.0 | (10, 10) | 540 |
| | | rep3 | (3, 2) | 71 | | | | |
| GFP negative | Fed Saline x3 | rep1 | (3, 2) | 70 | (3, 2) | 18.0 | (10, 10) | 300 |
| | | rep2 | (3, 2) | 66 | (2, 3) | 18.0 | (10, 10) | 500 |
| | | rep3 | (3, 2) | 70 | | | | |
| | Fas Saline x3 | rep1 | (3, 2) | 70 | (3, 2) | 18.0 | (10, 10) | 300 |
| | | rep2 | (3, 2) | 66 | (2, 3) | 18.0 | (10, 10) | 500 |
| | | rep3 | (3, 2) | 70 | | | | |
| | Fas Leptin x1 | rep1 | (3, 2) | 70 | (3, 2) | 18.0 | (10, 10) | 300 |
| | | rep2 | (3, 2) | 66 | (2, 3) | 18.0 | (10, 10) | 500 |
| | | rep3 | (3, 2) | 70 | | | | |
| | Fas Leptin x3 | rep1 | (3, 2) | 70 | (3, 2) | 18.0 | (10, 10) | 300 |
| | | rep2 | (3, 2) | 66 | (2, 3) | 18.0 | (10, 10) | 500 |
| | | rep3 | (3, 2) | 70 | | | | |

### Supplementary Table 4. Sequences of primers used for genotyping, cloning and qPCR and sgRNAs used for CRISPRi.

| | |
|---|---|
| Mouse genotyping for LepRcre and SUN1-sfGFP | |
| Lepr WT.F | GCCCTCATTAATCTAGTAATGTAGAT |
| Lepr WT.R | GCAAAAAAAGTAGTTAACCTATTCCT |
| Cre.F | CGATGCAACGAGTGATGAGG |
| Cre.R | GCATTGCTGTCACTTGGTCGT |
| INTACT.F | GCACTTGCTCTCCCAAAGTC |
| INTACT WT.R | CATAGTCTAACTCGCGACACTG |
| INTACT MT.R | GTTATGTAACGCGGAACTCC |
| Socs3 enhancer cloning | |
| Socs3_1.F | TGGCCTAACTGGCCGGTACCTCCCACACAGGATCAGCTTCCAAG |
| Socs3_1.R | TCTAGTGTCTAAGCTTAGCAGGTCCCAAGCTAGGTATGAG |
| Socs3_2.F | TGGCCTAACTGGCCGGTACCCTAAGTAAATAGTCACCGACCATCTG |
| Socs3_2.R | TCTAGTGTCTAAGCTTTTGCCTCTGGACCATTGCACCCAC |
| Socs3_3.F | TGGCCTAACTGGCCGGTACCGGCCATCTGGATTCAACAGGATCC |
| Socs3_3.R | TCTAGTGTCTAAGCTTCTCTGATCAAGATTTCTTGTCTGGG |
| sgRNA cloning. sgRNA sequences are underlined. | |
| Socs3_sgRNA_1.F | CCCTTGGAGAACCACCTTGTTGGTGACATTGTCACAGAAGTGGTTTAAGAGCTAAGCTGGAAACAGCA |
| Socs3_sgRNA_2.F | CCCTTGGAGAACCACCTTGTTGGAGGCTGATGTTCCTGTGAGGTTTAAGAGCTAAGCTGGAAACAGCA |
| pLG1.R | GATCCTAGTACTCGAGAAAAAAAGCACCGAC |
| qPCR | |
| Socs3.F | ACCTCGCAGATCCCTTGCACC |
| Socs3.R | TCTGCCTCCCTTCGGTGTTGG |
| Pgs1.F | GCTTCATCTCAGCCTTCTCAA |
| Pgs1.R | TTACACAGGAGGGATCAGGAA |
| Tha1.F | TAGCAGGATTGCCTAGTGTGC |
| Tha1.R | GCTTCTAATCCCCTAGCATGG |
